# T7 RNA polymerase-based gene expression from a transcriptionally silent rDNA spacer in the endosymbiont-harboring trypanosomatid *Angomonas deanei*

**DOI:** 10.1101/2025.01.30.635516

**Authors:** Lena Kröninger, Anay K. Maurya, Christian Stiebeling, Florian P. Stirba, Zio Kim, Eva C. M. Nowack

## Abstract

Eukaryotic life has been shaped fundamentally by the integration of bacterial endosymbionts. The trypanosomatid *Angomonas deanei* that contains a β-proteobacterial endosymbiont, represents an emerging model to elucidate initial steps in symbiont integration. Although the repertoire of genetic tools for *A. deanei* is growing, no conditional gene expression system is available yet, which would be key for the functional characterization of essential or expression of toxic proteins. Development of a conditional expression system based on endogenous RNA polymerase II (POLII) is hampered by the absence of information on transcription signals in *A. deanei* as well as the unusual genetic system used in the Trypanosomatidae that relies on read-through transcription. This mode of transcription can result in polar effects when manipulating expression of genes in their endogenous loci. Finally, only few resistance markers are available for *A. deanei* yet, restricting the number of genetic modifications that can be introduced into one strain. To increase the range of possible genetic manipulations in *A. deanei*, and in particular, build the base for a conditional expression system that does not interfere with the endogenous gene expression machinery, here we (i) implemented two new drug resistance markers, (ii) identified the spacer upstream of the rDNA array on chromosome 13 as transcriptionally silent genomic locus, and (iii) used this locus for engineering an ectopic expression system that depends on the T7 RNA polymerase expressed from the δ-amastin locus. We show that transgene expression in this system is independent of the activity of endogenous RNA polymerases, reaches expression levels similar to the previously described POLII-dependent expression from the γ-amastin locus, and can be applied for studying endosymbiosis. In sum, the new tools expand the possibilities for genetic manipulations of *A. deanei* and provide a solid base for the development of an ectopic conditional expression system.

## Introduction

The trypanosomatid *Angomonas deanei* is an emerging model system for endosymbiosis [1,2]. It harbors a β-proteobacterial endosymbiont of the Alcaligenaceae, which co-evolved with its host for 40-120 million years [3]. The endosymbiont supplies its host with essential metabolites and co-factors, underwent pronounced genome reduction [4], and divides strictly synchronously with its host cell [5], indicating an advanced level of endosymbiont integration. However, information on the exact molecular mechanisms of host-symbiont communication and control is scarce.

A few genetic tools have been developed for the asexual diploid *A. deanei*. Protocols available allow to efficiently generate heterozygous and homozygous knock-out mutants by one or two rounds of homologous recombination, respectively [6,7], or by CRISPR/Cas-based technology [8]. Furthermore, two amastin loci, called the δ-amastin and γ-amastin locus, coding for highly expressed surface glycoproteins that are dispensable for the organism under standard growth conditions in the laboratory, were successfully hijacked for high level transgene expression [6]. Expression of fluorescent fusion proteins from these loci has been instrumental for the identification of a number of <u>e</u>ndosymbiont targeted host <u>p</u>roteins (short ETPs) that are assumed to be involved in host/symbiont interaction [6,7]. However, inducible gene expression systems that would be important for the expression of toxic gene products and the development of conditional gene knock-outs of essential genes are not available yet, and although a protocol for RNA interference-based gene knock-downs has been published [9], so far, it has not been reproducible in other laboratories. These limitations call for the development of further genetic tools.

Trypanosomatids use an uncommon gene expression system. They produce long polycistronic mRNA strands from unidirectional gene clusters containing dozens of genes [10,11]. During mRNA maturation, polycistronic mRNAs are processed by coupled trans-splicing and polyadenylation reactions resulting in mature monocistronic mRNAs, that carry at their 5’ ends a common spliced leader (a capped 39-nt RNA molecule) [10,12]. Regulation of gene expression occurs on the mRNA level by stabilizing elements in the 5’ and 3’ untranslated regions (UTRs) directly influencing gene expression [10]. Consequently, transgenes that are inserted into any expressed genomic locus are transcribed by read-through, but their expression cannot be conditionally modulated or interrupted, e.g. using operator/repressor pairs, without affecting expression of downstream genes. This makes the development of inducible gene expression systems challenging.

Due to their medical relevance, other trypanosomatids such as *Trypanosoma* spp. and *Leishmania* spp. that do not carry an endosymbiont, are far better studied and advanced genetic tools are available for them [13–15]. These tools include conditional gene expression systems, that make use of transcriptional regulators, such as the tetracycline-inducible repressor (TetR) [16] or the cumate repressor (CymR) [17]. A common strategy to prevent interference with the endogenous gene expression machinery in these systems, is ectopic expression from transcriptionally silent genomic loci. Such silent loci have been found, for example, in transcriptionally inactive mini-chromosomes in *Trypanosoma brucei* [18] (which do not exist in *A. deanei*) or upstream of the promoter for the 18S rRNA gene (hereafter referred to as the rDNA spacer) [19,20]. In the latter insertion site, the inducible gene is inserted oriented in the opposite direction to that of the 18S rDNA [21].

In *T. brucei*, some promoter sequences such as the procyclic acidic repetitive protein (PARP) promoter have been described and used to establish conditional expression systems based on the endogenous POLII [19,21]. In *A. deanei*, however, virtually nothing is known about transcription initiation. Other expression systems in trypanosomatids make use of the exogenous bacteriophage T7 RNA polymerase (T7RNAP) [16,22–25]. Advantages of the T7RNAP are not only that it is a single subunit enzyme that can be produced heterologously in a wide variety of organisms but also that promoter and terminator sequences are well-characterized and comparably short [26]. Moreover, the enzyme shows an extraordinary high activity, beyond the activity of endogenous RNA polymerases, in some bacteria such as *Escherichia coli* [27]. In *T. brucei,* T7 RNAP-dependent gene expression exceeded POLII-dependent expression with a PARP promoter by 5-fold [22].

To increase the experimental versatility of *A. deanei* as an endosymbiosis model, here we describe a transcriptionally silent locus on its nuclear genome. We demonstrate that this locus can be used for T7RNAP-driven expression of transgenes. We compare expression strength of this newly introduced system to the previously described expression system from the amastin loci and implement blasticidin and nourseothricin resistance genes as new selectable markers in *A. deanei*. Together the new protocols contribute to the growing genetic toolbox that supports the establishment of *A. deanei* as a versatile endosymbiosis model.

## Results

### Identification of potentially silent genetic loci as integration site for transgenes in *A. deanei*

To identify potentially silent genomic loci in *A. deanei* that could be used for transgene expression by an exogenous transcription machinery, we searched the *A. deanei* nuclear genome for ribosomal repeats and found seven copies on different chromosomes (**Fig 1**). Putative protein-coding sequences upstream of the ribosomal arrays were identified using the auto-annotation tool of Benchling (https://www.benchling.com) and comparisons to the curated annotations in the TriTryp Database (https://tritrypdb.org). The first coding sequence upstream of the ribosomal arrays was found 3.2 kbp away from the 18S rDNA (on chromosome 25) suggesting that these intergenic spacers are indeed non-coding. All seven rDNA spacers show high sequence similarity in the first 1.5-2 kbp upstream of the ribosomal arrays, but then diverge with increasing distance from the ribosomal arrays (see alignment of the first 4000 bp upstream of the ribosomal array in **S1 Appendix**).

**Fig 1.**
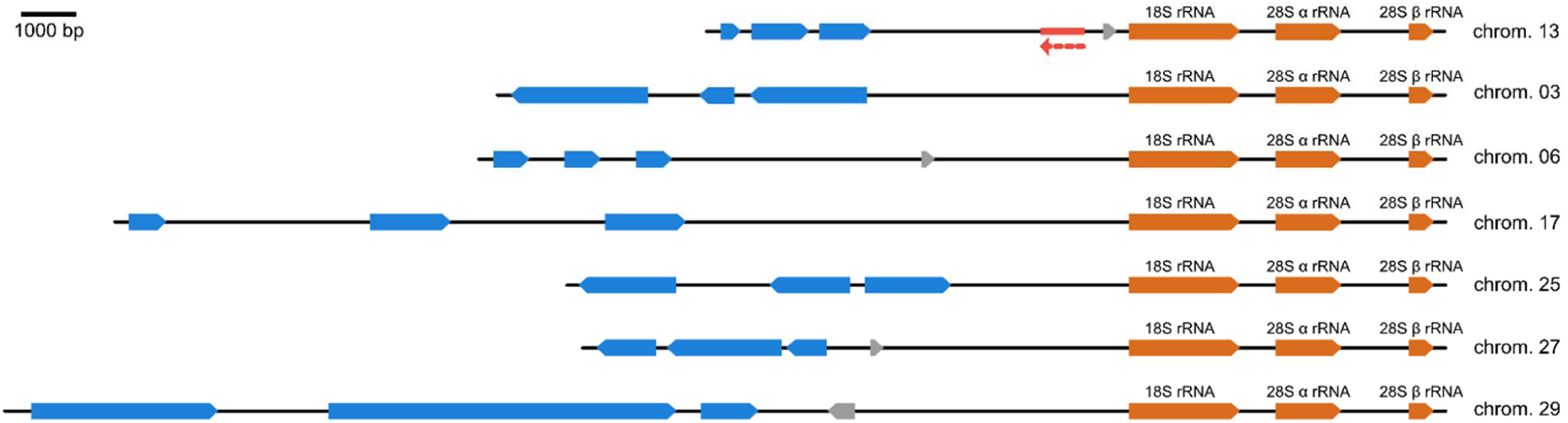
Potentially silent loci upstream of the ribosomal repeats in *A. deanei*. The highly conserved ribosomal repeats were found on seven different chromosomes (chrom.) and are marked in orange. Putative protein-coding sequences upstream of the ribosomal repeats are shown in blue. Open reading frames marked in grey did not show homologs in other trypanosomatids (using BLASTp against the NCBI non-redundant database) and are unlikely to represent true protein-coding genes. The targeted insertion site on chromosome 13 is marked in red and transgene orientation is indicated by the broken red arrow.

### Heterologous expression of the T7RNAP in *A. deanei*

To implement an RNA polymerase in *A. deanei* that works independently of the endogenous expression machinery, we inserted the T7RNAP coding sequence into the δ-amastin locus. For selection, the T7RNAP coding sequence was linked upstream via the intergenic region between GAPDH-I and II genes (GAPDH-IR, that contains signals for mRNA maturation and stabilization [6]) to the neomycin phosphotransferase gene (*neo^R^*) (**Fig 2A**). Transfectants were selected on neomycin (neo) and the clonal strain obtained (Adea319) was verified by PCR with primers binding outside of the integration site (**Fig 2B**). Production of the T7RNAP was verified by Western blot analysis (**Fig 2C**).

**Fig 2.**
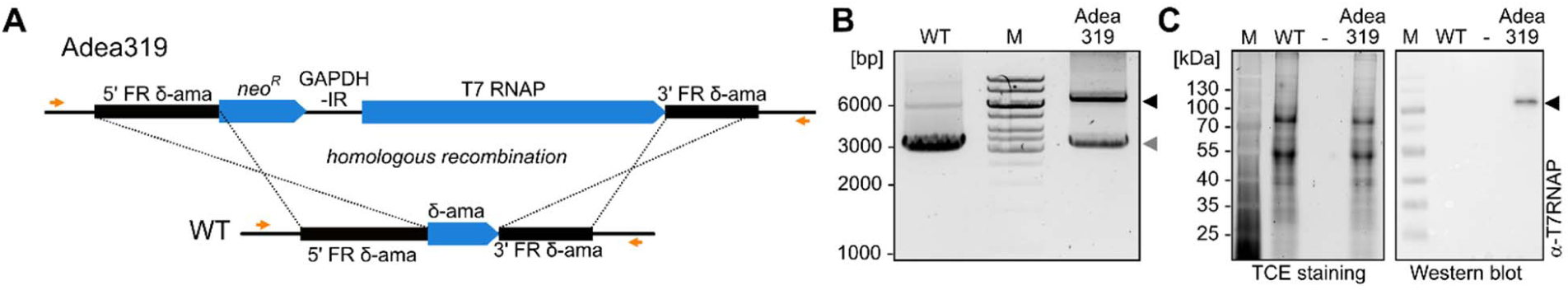
Construction and verification of the T7RNAP-producing strain Adea319. **(A)** Genomic map of Adea319 and the *A. deanei* wildtype (WT) in the δ-amastin (δ-ama) locus. Primers 49+545, that were used for the verification of the insertion of the T7RNAP gene-containing construct, are marked in orange. **(B)** Verification of the integration of the T7RNAP expression cassette in Adea319 by PCR. The WT served as a control and showed only the δ-amastin WT locus (3.3 kbp, grey arrowhead) plus a weak unspecific band at 6.0 kbp. The recombinant locus in Adea319 additionally yielded the predicted product at 6.2 kbp (black arrowhead). **(C)** Verification of production of the ∼100 kDa T7RNAP (black arrowhead) by Western blot using an α-T7RNAP primary antibody (right picture). Trichloroethanol (TCE) staining of the SDS-polyacrylamid gel electrophoresis (PAGE) before blotting confirmed equal loading of the WT and Adea319 raw extract samples (left picture).

### Implementation of new antibiotic resistance markers for *A. deanei*

So far, only a limited set of antibiotic resistance markers has been established for *A. deanei* strain ATCC PRA-265 [6,7], namely *neo^R^*, the hygromycin phosphotransferase (*hyg^R^)*, and the phleomycin-binding protein (*phleo^R^)*. To increase the number of resistance markers, we tested blasticidin (blast) and nourseothricin (nours) and their resistance genes blast deaminase (*blast^R^*) and nours acetyltransferase (*nours^R^*), respectively, as potential selection markers in *A. deanei*. To this end, we generated two strains, Adea373 and Adea384, expressing *blast^R^* (alone) and *nours^R^* (in combination with *neo^R^*) from the δ-amastin locus and compared their growth with the WT on various concentrations of the respective antibiotics. Whereas growth of the WT was heavily affected at blast concentrations ≥75 µg/ml and nours concentrations ≥200 µg/ml, Adea373 and Adea384 were resistant to the respective antibiotics, and grew up to concentrations of 200 µg/ml blast or 400 µg/ml nours, respectively, without noticeable growth defects (**Fig 3**). Subsequently, 75 µg/ml blast or 200 µg/ml nours were used for mutant selection.

**Fig 3.**
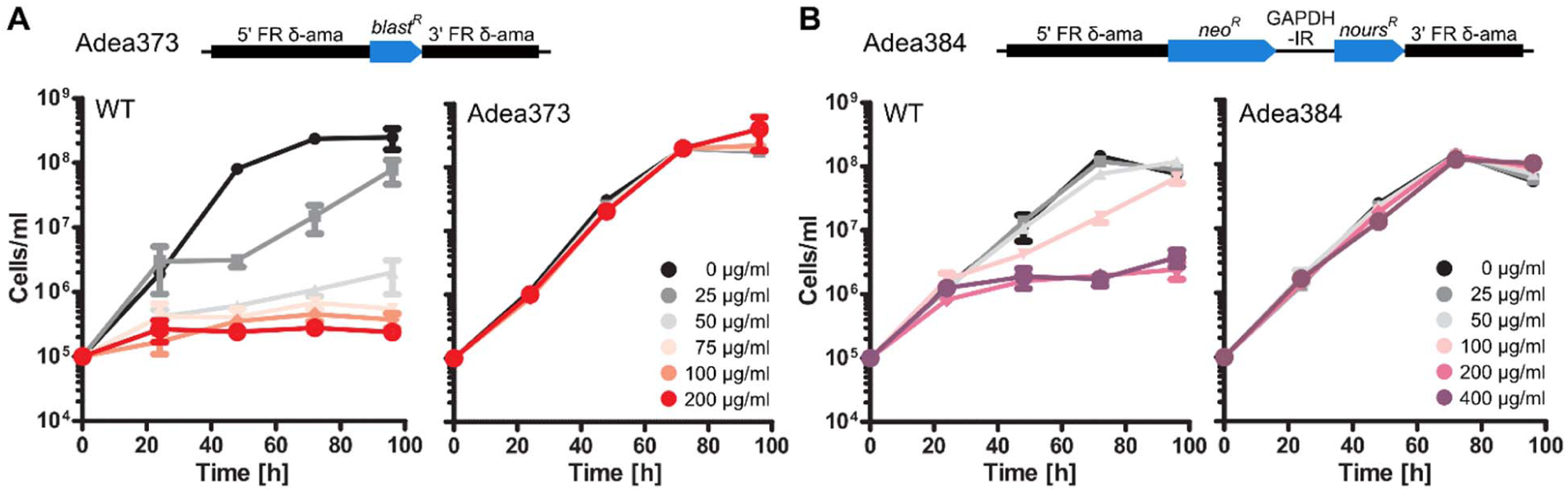
Susceptibility of *A. deanei* WT to blasticidin (blast) and nourseothricin (nours) compared to susceptibility of Adea373 to blast and Adea384 to nours, respectively. **(A)** Genomic map of Adea373 and growth of *A. deanei* WT and Adea373 on increasing blast concentrations over 4 days. **(B)** Genomic map of Adea384 and growth of *A. deanei* WT and Adea384 on increasing nours concentrations over 4 days. Blast (A) and nours (B) concentrations are color coded as indicated in the graphs. The graphs display mean values and standard deviations from 3 biological replicates.

### T7RNAP-driven expression from the rDNA spacer

Next, we aimed for the insertion of a transgene with T7 transcription signals into the rDNA spacer on chromosome 13 (chr13 rDNA spacer, see **Fig 1**) to test functionality of the T7RNAP. As selection marker and reporter cassette we made use of *blast^R^* flanked by sections of intergenic regions that trigger trans-splicing and polyadenylation during mRNA maturation, namely a 252 nt-long fragment of the intergenic region upstream of the δ-amastin gene (“spliced leader donor”, SLD) as 5’ flank and the complete GAPDH-IR as 3’ flank (**Fig 4A**). The T7 promoter and the T7 terminator were added at the 5’ and 3’ ends of this cassette, respectively. Finally, 600 bp-long sequences flanking the genomic target locus (see **S1 Appendix**) were added upstream and downstream of the T7 transcription signals for homologous recombination. Next, the T7RNAP-producing strain, Adea319, was transfected with this *blast^R^* integration cassette and blast was added to the transfectants 6 h post transfection.

**Fig 4.**
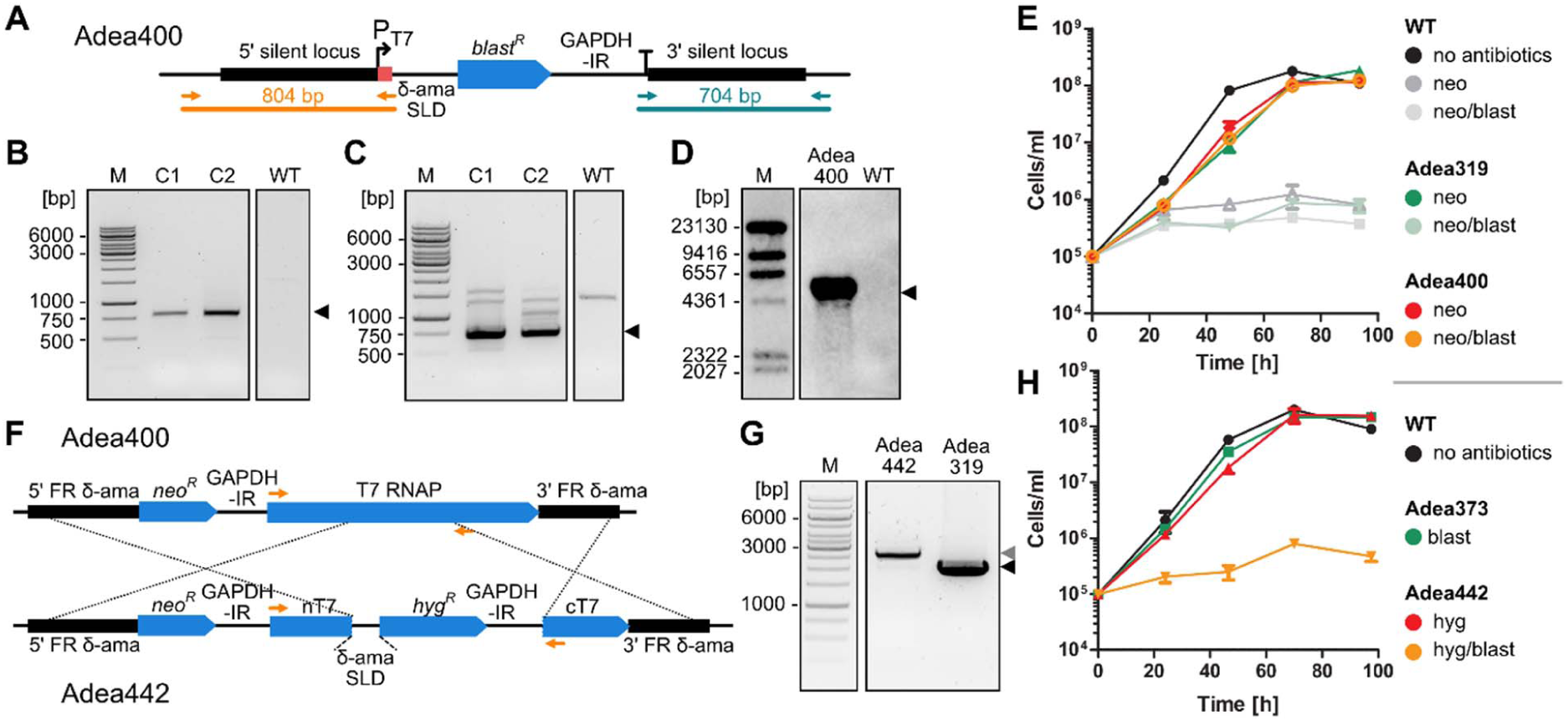
T7RNAP drives *blast^R^* expression from the rDNA spacer. **(A)** Genomic map of the chr13 rDNA spacer of Adea400. Primer pairs 2211+2748 and 2753+2210 that were used for the verification of the insertion of the *blast^R^*-containing construct and expected products are shown in orange and blue-green. **(B, C)** Verification of integration of the *blast^R^*-containing construct into the targeted rDNA spacer by PCR (B, orange primer pair 2211+2748 and C, blue-green primer pair 2753+2210). The WT served as a control and showed no (B) or a faint unspecific product (C), whereas samples of two individual clones (C1 and C2) yielded the expected product at ∼800 bp (B) and ∼700 bp (C). **(D)** Verification of insertion into the chr13 rDNA spacer in Adea400 by Southern blot. The *blast^R^*-targeting probe yielded only a single band of the expected size at 4746 bp for Adea400 but not for the WT control (see **S1 Table** for expected sizes and **S1 Figure** for an additional Southern blot on BamHI-digested DNA). **(E)** Growth of *A. deanei* WT, Adea319, and Adea400 on neo and neo/blast over 4 days. **(F)** Knock-out strategy and genomic map of Adea442 depicting the interruption of the T7RNAP-encoding gene by a *hyg^R^*-containing cassette and partial removal of the gene with only N- and C-terminal domains (nT7/cT7) remaining. **(G)** Verification of the successful T7RNAP knock-out by PCR using primers 2045+2966 (labelled in orange in the genomic map). Adea319 with functional T7RNAP gene (black arrowhead), Adea442 with successfully disrupted T7RNAP gene (grey arrowhead). **(H)** Growth of *A. deanei* WT, Adea373, and the T7RNAP knock-out strain Adea442 on hyg, blast, and hyg/blast over 4 days. In (E) and (H) mean values and standard deviations from 3 biological replicates are displayed.

Recombinant clones recovered after 5-7 days suggesting that *blast^R^* is expressed in sufficient amounts to render the cell resistant. As mentioned above, the rDNA spacers on different chromosomes are highly similar, so that i) insertion into a non-targeted locus and ii) multiple insertions into several rDNA spacers could not be excluded. To explore these issues, first we verified the insertion of the *blast^R^*-containing construct into an rDNA spacer by PCR (**Fig 4B, C**). Then, we identified BclI and SpeI as restriction enzymes that allow to distinguish between insertions into the rDNA spacers on different chromosomes by Southern blot analysis due to restriction length polymorphisms (see **S1 Table**). Southern blot analysis of Adea400 with BclI and SpeI yielded a single band of 4746 bp, indicative of an insertion specifically into the chr13 rDNA spacer (**Fig 4D**). Lastly, growth experiments (**Fig 4E**) revealed that the doubling time of Adea400 on blast+neo (6.9 ± 0.2 h) and the T7RNAP-producing strain Adea319 on neo alone (6.9 ± 0.2 h) were identical. And both strains grew only slightly slower that the WT without antibiotics (doubling time of 6.6 ± 0.1 h).

To exclude that the endogenous transcription machinery of *A. deanei* is responsible for *blast^R^* expression from the rDNA spacer, we generated a strain derived from Adea400 (named Adea442), in which the T7RNAP-encoding gene was interrupted by a *hyg^R^* cassette as well as partially removed (**Fig 4F**). We first verified the successful insertion into the target locus by PCR (**Fig 4G**) and then tested the verified strain for blast resistance. On hyg alone, Adea442 showed similar growth rates (doubling time 6.6 ± 0.4 h) as *A. deanei* WT without antibiotics (doubling time 6.1 ± 0.3 h) and the blast-resistant strain Adea373 on blast (doubling time 6.4 ± 0.1 h) (**Fig 4H**). However, no growth of Adea442 was observed on blast. This result clearly demonstrates that *blast^R^* expression from the chr13 rDNA spacer is controlled by the T7RNAP system and not the endogenous transcription machinery.

### Analysis of gene expression strength

To compare the expression strength between the T7RNAP-dependent system and POLII-dependent expression from the amastin loci, we generated a strain that expresses the red fluorescent protein mScarlet fused to ETP1 (Adea456) from the chr13 rDNA spacer. ETP1 is one of the host-encoded ETPs and localizes at the periphery of the endosymbiont [7]. The coding sequence for the fusion protein was part of an integration cassette containing *blast^R^* as selection marker (**Fig 5A**).

**Fig 5.**
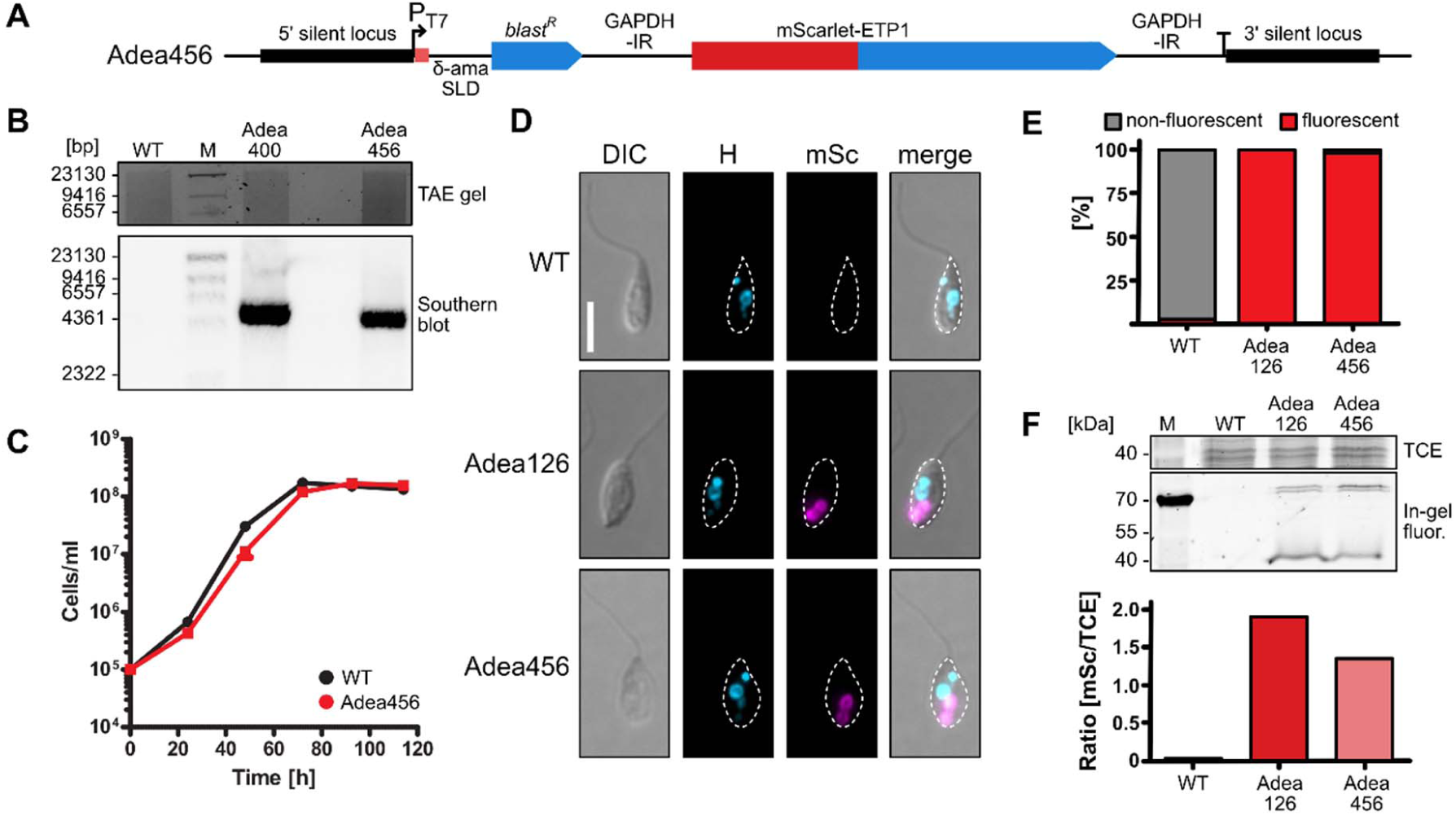
Quantification of T7RNAP-dependent protein expression from the rDNA spacer. **(A)** Genomic map of the chr13 rDNA spacer of Adea456 containing the gene encoding mScarlet-ETP1. **(B)** Verification of insertion in the chr13 rDNA spacer in Adea456 by Southern blot with BclI and SpeI. The WT and blast resistant strain Adea400 served as controls. The *blast^R^*-targeting probe yielded single bands of the expected sizes for Adea400 (4746 bp) and Adea456 (4345 bp) but not for the WT (see **S1 Table** for expected sizes). **(C)** Growth of *A. deanei* WT without antibiotics and Adea456 on neo/blast over 4 days. **(D)** Epifluorescence microscopic analysis of Adea126 and Adea456 expressing mScarlet-ETP1. Cell shape is indicated by white dashed lines. DIC, differential interference contrast; H, blue channel (visualizing Hoechst 33342); mSc, red channel (visualizing mScarlet). Scale bar represents 5 µm. **(E)** Quantification of fluorescent and non-fluorescent cells of the WT, Adea126, and Adea456 based on epifluorescence microscopy pictures. For each cell line at least 250 cells were analyzed. **(F)** In-gel fluorescence assay detecting red fluorescence in raw extracts of the WT, Adea126, and Adea456. TCE, TCE-stained total protein; In-gel fluor., in-gel fluorescence; M, marker. Note that the assay conditions result in incomplete protein denaturation resulting in several bands for mScarlet-ETP1 (69 kDa).

First, targeted insertion of the integration cassette into the chr13 rDNA spacer in Adea456 was verified by Southern blot (**Fig 5B** and **S1 Figure**). While the DNA load seemed to be slightly lower in Adea400 compared to Adea456 (see gel picture), the Southern blot signal was more pronounced in Adea400. Hence, it is possible that the integration cassette inserted twice (in both alleles) in Adea400 but only once in Adea456. Growth of Adea456 (doubling time of 6.6 ± 0.1 h) was only slightly slower than in the WT (6.3 ± 0.1 h) (**Fig 5C**). Next, the expression level of mScarlet-ETP1 was analyzed by an in-gel fluorescence assay and homogeneity of expression by epifluorescence microscopy. The WT and Adea126, which produces mScarlet-ETP1 from the γ-amastin locus [7], served as references. Microscopically, Adea456 and Adea126 showed comparable mScarlet fluorescence patterns (**Fig 5D** and **S2 Figure**) with the endosymbiont being labelled in its periphery in 98 and 100 % of the cells, respectively (**Fig 5E**). Using the same microscope settings, in only ∼1 % of the WT cells a weak autofluorescent signal was recorded. The in-gel mScarlet fluorescence intensity (normalized to TCE-stained total protein) for Adea456 was approximately 20 % weaker than for Adea126 but still in a similar range (**Fig 5F**).

## Discussion

Bacterial endosymbionts are widespread across eukaryotes and profoundly impact physiology and evolution of their hosts [28–30]. Although the genomes of countless bacterial endosymbionts have been sequenced and hence, metabolic complementation between host and endosymbiont are quite well understood in many systems, our understanding of the molecular mechanisms that enable metabolic complementation, signaling, and exertion of control between the symbiotic partners lacks far behind. A central problem for dissecting these questions is that endosymbiotic systems that readily grow in the laboratory and can be efficiently genetically modified are scarce. With the current work we contribute to further develop *A. deanei* into such a much sought after, versatile model for the dissection of host/endosymbiont interactions. To this end, we implemented new resistance markers, identified a transcriptionally silent genomic locus, and, most importantly, used this locus to develop an ectopic gene expression system that is independent of the endogenous transcription machinery.

As new selectable markers we introduced *nours^R^* and *blast^R^* and determined by titration 75 µg/ml blast or 200 µg/ml nours as selective concentrations that efficiently kill the WT cells whereas resistance cassette-carrying transformants are unaffected (**Fig 3**). These selection markers are commonly used in other trypanosomatids but effective selective concentrations vary between strains. E.g. inhibitory concentrations of 0.7-10 µg/ml blast [31,32] and 2-50 µg/ml nours [31,33] are reported for *Leishmania* sp.; and 2-10 µg/ml blast [32,34] and 500 µg/ml nours [35] for *Trypanosoma* sp.. Together with the previously implemented *neo^R^*, *hyg^R^*, and *phleo^R^*, these additional selectable markers allow to introduce up to five selectable cassettes into the same *A. deanei* strain now.

As an insertion site for ectopic gene expression, we chose the chr13 rDNA spacer into which expression cassettes were inserted in reverse orientation with respect to the rDNA array. Inactivity of endogenous RNA polymerases in this locus was evident by the sensitivity of strain Adea442, that carries a *blast^R^* T7 expression cassette in the chr13 rDNA spacer but no functional T7RNAP gene, to selective concentrations of blast (**Fig 4H**). Although the exact RNA polymerase I promoter that is needed for the production of rRNAs from the rDNA arrays, is not known in *A. deanei*, studies from *T. brucei* suggest that the promoter and the transcriptional start are ∼1.2 kbp upstream of the 18S rDNA [20,36,37]. If the promoter is in the same position in *A. deanei*, it would be within the 5’ flank that was used for homologous recombination, resulting in insertion of the cassette upstream of this putative promoter region (see **S1 Appendix**). Hence, it is unlikely that integration of expression cassettes into this site interferes with transcription of the rDNA array downstream and our growth analyses demonstrate that even in strain Adea400, in which insertion of the expression cassette apparently occurred in both copies of chr13, growth is not impaired compared to its parental strain Adea319 which expresses only T7RNAP but carries no additional expression cassettes in the chr13 rDNA spacer (**Fig 4E**). Although the 5’ flank that we used for homologous recombination showed a high degree of sequence similarities between the rDNA spacers of the seven different chromosomes (see **Fig 1** and **S1 Appendix**), in all clones that we obtained from transfections with insertion cassettes that carry 600 bp flanks amplified from the chr13 rDNA spacer, homologous recombination occurred into the chr13 rDNA spacer. Hence, the 3’ flank that shows more sequence divergence between different chromosomes, apparently was sufficient to guide insertion of the cassette specifically into the targeted integration site.

As an exogenous RNA polymerase for the ectopic gene expression system, we relied on the T7RNAP expressed from the δ-amastin locus. Although the δ-amastin locus is up to this point the strongest expression locus for transgenes known in *A. deanei a*nd ranges on rank 35 of the most highly expressed genes in *A. denaei* [6], T7RNAP expression did not result in pronounced toxicity as evidenced by the only very mild growth reduction of T7RNAP-expressing cell lines compared to WT (**Fig 4E**). In *E. coli,* high T7RNAP levels can be toxic [38] and similar effects were observed in *T. brucei*. In the latter, initial attempts to generate transgenic lines that express the T7RNAP from the highly expressed PARP locus, were repeatedly unsuccessful [39]. This obstacle was solved by placing the T7RNAP-encoding gene into the weaker β-tubulin expression locus [19], suggesting that high T7RNAP concentrations are lethal.

At the same time, the T7RNAP levels obtained by expression from the δ-amastin locus resulted in high levels of T7RNAP-driven expression of transgenes that are flanked by T7 transcription signals. T7RNAP-dependant expression of *blast^R^* (strain Adea400) rendered cells resistant to the same selective blast concentration as cells with POLII-dependent expression of *blast^R^* from the δ-amastin locus (strain Adea373) (compare **Figs 3A** and **4E**). Furthermore, T7RNAP-dependent expression of mScarlet-ETP1 (strain Adea456), resulted in protein levels that were well detectable by epifluorescence microscopy (**Fig 5D**) and only 20 % lower than those that were obtained when expressing the same fusion protein POLII-dependently from the γ-amastin locus (strain Adea126) (**Fig 5F**). These results highlight the new expression system as a suitable tool for investigating symbiosis-related proteins such as the ETPs.

Finally, the new ectopic expression system and resistance markers now provide the basis for the development of advanced gene expression systems, in which the ectopic expression of a gene of interest can be modulated by transcriptional regulators, such as TetR (in its ON or OFF variant [21,40,41]) or CymR [42] without triggering polar effects caused by interference with the endogenous gene expression machinery. Conditional expression would allow to study the effect of the expression of toxic proteins. Furthermore, conditional ectopic expression of a copy of a gene of interest in its null mutant background would enable conditional gene knock-out strategies that would leverage the dissection of cellular functions of essential gene products. Expression strength of this system can be modulated, if needed, by (i) taking advantage of known T7 promoter variants with decreased [43,44] or increased activity [45], (ii) increasing the copy number of the transgene integrated into the same or different rDNA spacers or (iii) by integrating the T7 expression cassettes into episomal elements (e.g. plasmids) which is routinely used, for example, in *L. tarentolae* [46] and *T. brucei* [39].

## Materials and methods

### Generation of transgenic cell lines

*A. deanei* (ATCC PRA-265) was grown in brain heart infusion (BHI, Sigma Aldrich) medium supplemented with 10 μg/ml of hemin (Sigma Alderich) at 28 °C as described before [6]. Recombinant strains were generated by transfection with specific integration cassettes that had been excised from their storage vectors prior to transfection (see **S2 Table** and **S3 Table**). For the excision of the integration cassettes EcoRI or EcoRV/EcoRI (both New England Biolabs) were used. Transfections were performed as described before [7]. After transfection and recovery, cells were mixed with specific antibiotics for selection [neomycin/G418 (Sigma-Aldrich) at 500 µg/ml, hygromycin B (Invivogen) at 500 µg/ml, blasticidin S (Invivogen) at 75 µg/ml, and nourseothricin (Carl Roth) at 200 µg/ml] before dilution and distribution over a 96-well plate [7]. Cells typically recovered after 5-9 days, and correct insertion of the transgenes was verified by PCR and/or Southern blot analysis. A list of all strains used in this study is given in **S3 Table**.

### Growth analyses

For the analysis of the growth behavior of different *A. deanei* strains, precultures were grown to mid-exponential phase (approximately 1-5 × 10^7^ cells/ml) with all selective drugs that would allow growth of the specific strains. Then, three replicates were prepared with 5 ml BHI plus hemin and antibiotics as indicated and inoculated to an initial cell density of 1 × 10^5^ cells/ml. The tubes were incubated at 28 °C without agitation and the cell density was measured every 20-24 h using the MultiSizer 4e (Beckman Coulter).

### Analysis of homologous recombination by Southern blot

Southern blot was performed as described previously [7]. Each DNA sample contained 7 µg of purified DNA. The DNA was digested using restriction enzymes BclI and SpeI or only BamHI (New England Biolabs). A 393 bp DIG-labelled probe against the blasticidin resistance gene (*blast^R^*) was generated by PCR using primers 3174 and 3175 (see **S3 Table**) and the DIG DNA Labelling Mixture (Roche) according to the manufacturer’s instructions. Hybridization was conducted at a temperature of 48 °C.

### Construction of plasmids

All plasmids for this study were products of Gibson [47] or Golden Gate [48] assemblies as described previously [7]. The cloning fragments, which were needed for the assemblies, were generated by PCR using the Phusion polymerase (New England Biolabs) and the primers and templates are listed in **S2 Table**. The correct sizes of cloning fragments were verified by electrophoresis. PCR products matching the expected sizes were excised from the gel and purified using the Monarch DNA Gel Extraction kit (New England Biolabs). The concentration of the DNA was determined using the NanoPhotometer NP80 (Implen). The molar concentration of the insert(s) usually exceeded the vector backbone concentration by a factor of 3. After the assembly, 1 µl of DpnI was added to each assembly mix to remove residual PCR template present in the mix. Then, *E. coli* TOP10 cells were transformed with the assembly mix and recombinant cells selected on lysogeny broth (LB) plates containing 100 µg/ml ampicillin. Recovered clones were inoculated in 5 ml LB liquid medium with ampicillin (see above) and used for plasmid purification (Monarch Plasmid Miniprep kit, New England Biolabs). Plasmids were verified by control digests and sequencing (Microsynth).

### Epifluorescence microscopy

Microscopy slides were prepared from mid-exponential cultures (approximately 1-5 × 10^7^ cells/ml) as described before [7]. Images were taken with the Axio Imager A.2 equipped with the AxioCam MRm and an Illuminator HXP 120 V (all instruments from Zeiss) and processed with Zen Blue v2.5 (Zeiss) as described previously [7]. *A. deanei* WT cells served as negative control and its autofluorescence was used to define a suitable fluorescence threshold. For post-processing of the pictures Fiji [49] was used.

### Western blot

For the detection of specific proteins, we grew 10 ml cultures of the relevant strains to mid-exponential phase, harvested the cells by centrifugation, and precipitated the proteins using trichloroacetic acid [7]. The precipitate was resuspended in urea buffer (8 M urea, 100 mM Tris, 5.25 mM EDTA, pH 8.6) and the protein concentration was determined using Pierce 660nm Protein Assay Reagent (ThermoFisher) and the FLUOstar Omega Plate Reader (BMG Labtech). A bovine serum albumin (BSA) standard curved served as reference. For each sample, 25 µg protein were mixed with Lämmli sample buffer (final concentrations: 63 mM Tris-HCl, pH 6.8, 10 mM dithiothreitol, 10% v/v glycerol, 2% SDS w/v, and 0.0025% w/v bromophenol blue), incubated for 5 min at 95 °C, and loaded onto a 10 % SDS polyacrylamid gel containing 0.5 % TCE. PAGE and protein blot to a PVDF membrane (Amersham Hybond, 0.45 nm, GE HealthCare Life Science) were performed as described previously [7]. After blotting, proteins were detected immunologically. In brief, the membrane was blocked with 5 % w/v milk powder for 1 h at room temperature (or 24 h at 4 °C) and washed three times for 5 min with 1x PBST (phosphate-buffered saline including 0.05 % v/v Tween-20) before the primary antibody was added. For detection of T7RNAP, an α-T7RNAP antibody (29943-1-AP, Proteintech) was used in a 1:5000 dilution overnight at 4 °C. Before the secondary conjugate was added, the membrane was washed again three times for 5 min with 1x PBST. The α-T7RNAP antibody was detected using α-rabbit IgG HRP conjugate (7074, CellSignaling), which was applied for 1 h at room temperature followed by three more washing steps (see above). Chemiluminescence was started by the addition of SuperSignal West Pico Chemiluminescent Substrate (ThermoFisher) and it was recorded using the ChemiDoc MP (Biorad).

### In-gel fluorescence assay

For the detection of in-gel fluorescence of mScarlet, samples were prepared as described by Sanial et al. [50] with minor modifications. First, 15 ml cultures of the relevant strains were grown to mid-exponential phase, cells were harvested by centrifugation, washed twice using 1x PBS, and resuspended in 1 ml 1x PBS. Before cell lysis, cOmplete EDTA-free protease inhibitor tabs (Roche) and a few grains of DNase (Carl Roth) were added and the cells were lysed by sonication using the UP100H (Hielscher) with 5 x 18 pulses (cycle 0.8, amplitude ∼60 %, MS1 sonotrode). Cell debris and intact cells were removed by centrifugation (3000 rpm, 4 °C, 5 min) and the protein concentration of the supernatant was determined using the Pierce 660nm Protein Assay Reagent (ThermoFisher) and the FLUOstar Omega Plate Reader (BMG Labtech). 25 µg of protein were mixed with Lämmli sample buffer (final concentrations: 50 mM Tris-HCl, pH 6.8, 10% v/v glycerol, 2% SDS w/v, 2.5 mM EDTA, and 0.005% w/v bromophenol blue, plus 12.5 mM dithiothreitol), heated to 50 °C for 5 min and applied to a 12.5 % SDS polyacrylamid gel containing 0.5 % v/v TCE. The electrophoresis ran at 40 mA until the marker bands (Prestained Ladder, ThermoFisher) were fully resolved. Finally, mScarlet fluorescence was recorded with the ChemiDoc MP (Biorad) using the UV tray and the “Cy3” filter. Afterwards, loading of the gel was visualized using TCE fluorescence with the UV tray and the “Stain-free gel” setting of the ChemiDoc MP (Biorad). The intensities of the mScarlet and TCE bands were estimated with Fiji [49] and normalized by the TCE signals.

## Acknowledgements

This work was supported by the Deutsche Forschungsgemeinschaft (DFG, German Research Foundation) – SFB1535, Project ID 458090666.

## Author contributions

L.K. and E.C.M.N. designed the research. L.K., A.K.M, C.S., F.P.S., and Z.K. performed the research. L.K. and E.C.M.N. analyzed the data. E.C.M.N. supervised the research. L.K. and E.C.M.N. wrote the manuscript.

## Data Availability Statement

All relevant data are within the manuscript and its Supporting Information files.

## Supporting information

### Supplemental Figures

**S1 Figure.**
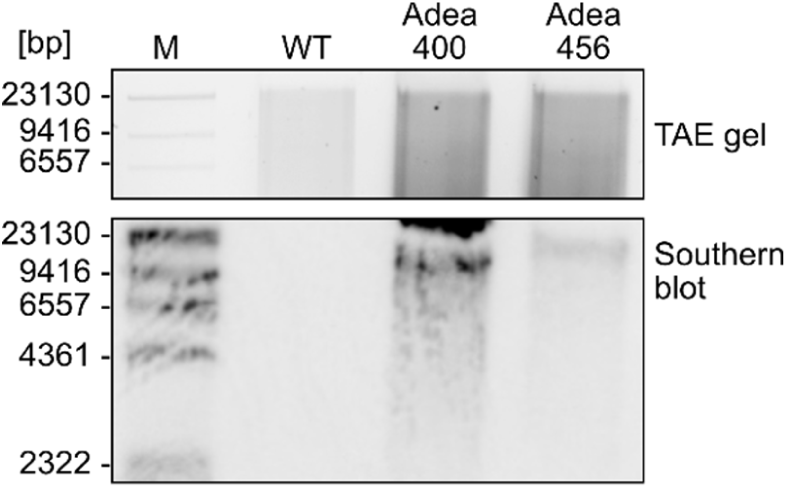
Verification of insertion in the chr13 rDNA spacer in Adea400 and Adea456 by Southern blot with BamHI. The WT served as control. The *blast^R^*-targeting probe yielded single bands of the expected sizes for Adea400 (16252 bp) and Adea456 (18519 bp) but not for the WT (see **S1 Table** for expected sizes).

**S2 Figure.**
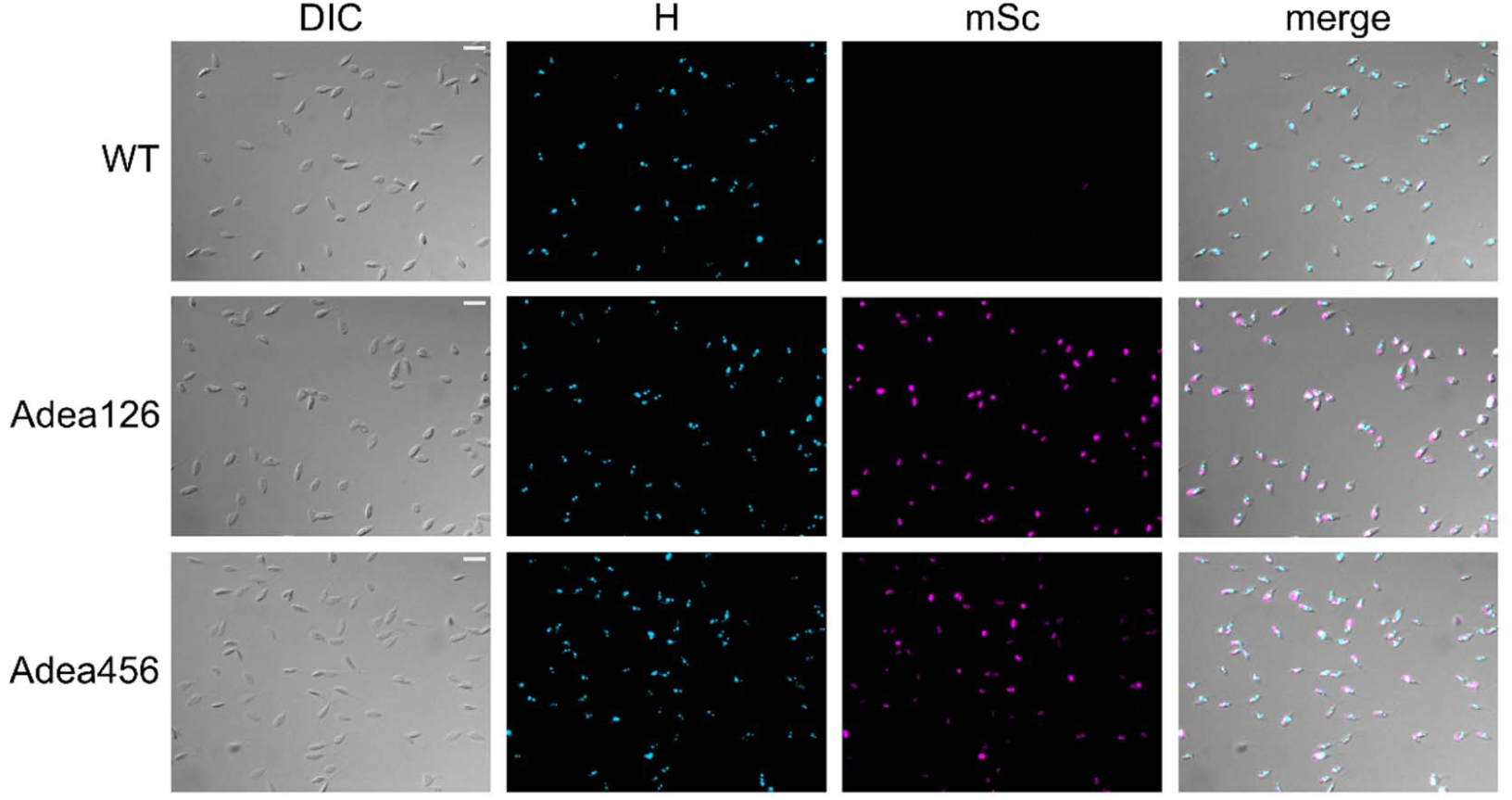
Epifluorescence microscopic analysis of Adea126 and Adea456 expressing mScarlet-ETP1. DIC, differential interference contrast; H, blue channel (visualizing Hoechst 33342-stained DNA); mSc, red channel (visualizing mScarlet). Scale bar represents 10 µm.

### Supplemental Tables

**S1 Table.** Expected Southern blot fragments for the verification of Adea400 and Adea456. For the identification of the rDNA spacer, in which transgenes had been inserted, genomic DNA of the relevant strains was digested using a combination of BclI and SpeI or BamHI alone.

| Strain | Length of expected band [bp] |  |  |  |
| --- | --- | --- | --- | --- |
|  | Adea400 |  | Adea456 |  |
|  | SpeI+BclI | BamHI | SpeI+BclI | BamHI |
| 13 | 4,746 | 16,252 | 4,345 | 18,519 |
| 3 | 8,142 | 4,535 | 4,332 | 6,802 |
| 6 | >30,000 | 5,520 | 4,353 | 7,787 |
| 17 | 7,926 | 5,515 | 4,347 | 7,782 |
| 25 | 4,943 | 7,453 | 4,355 | 9,720 |
| 27 | 10,999 | 5,525 | 4,356 | 7,792 |
| 29 | 14,036 | 8,131 | 4,356 | 10,398 |

## Other Supplemental Information

**S1 Appendix.**
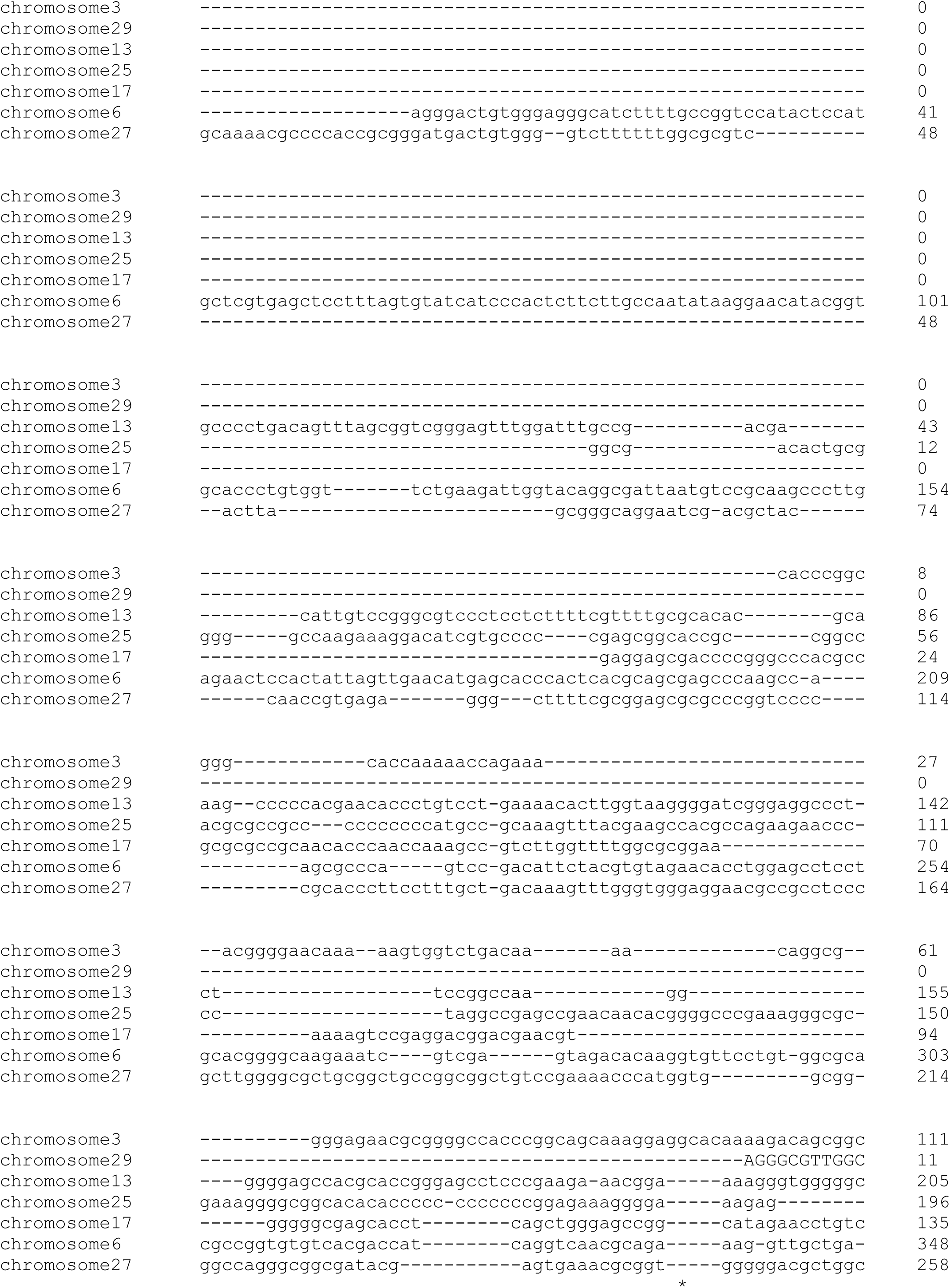
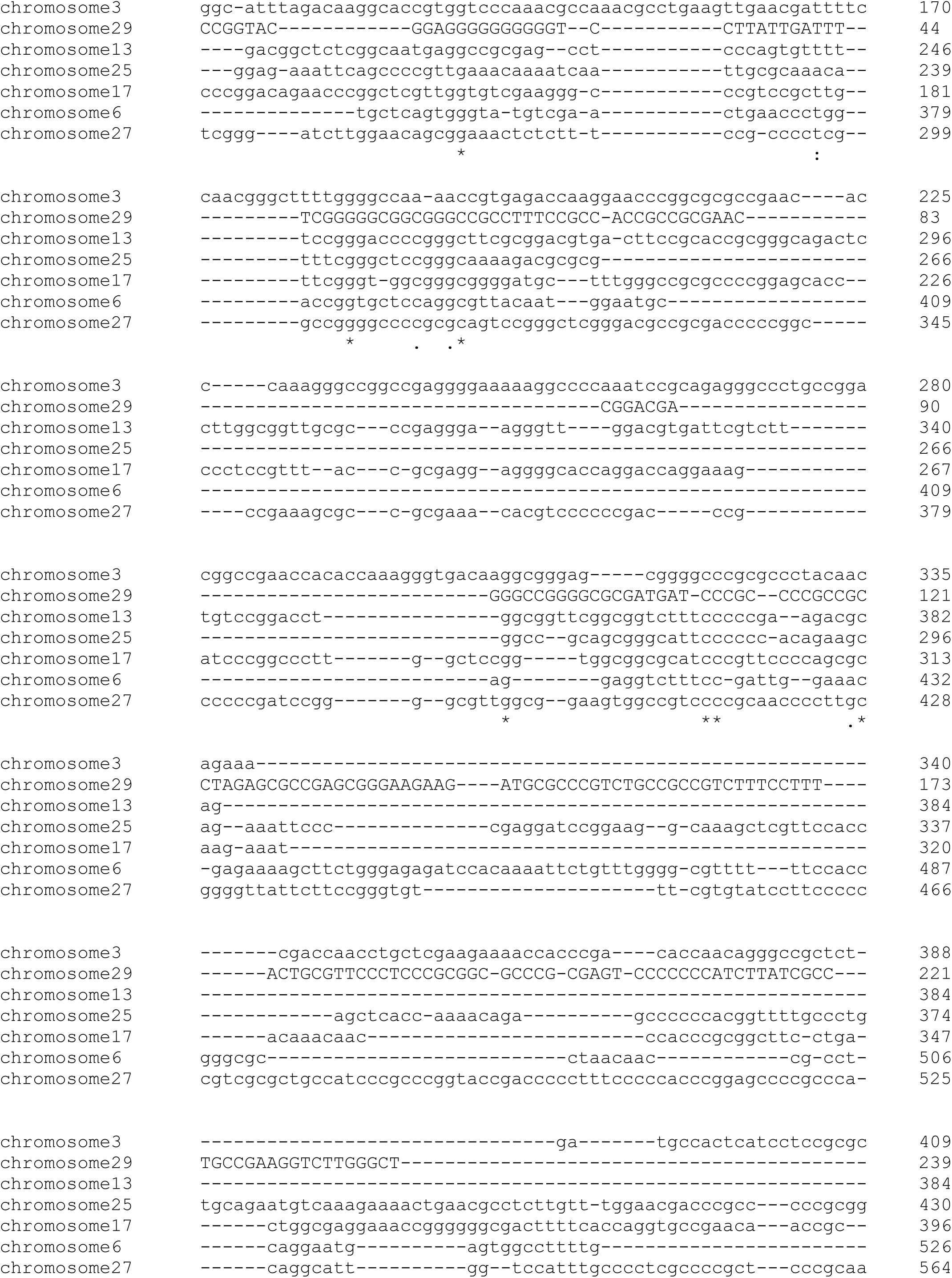
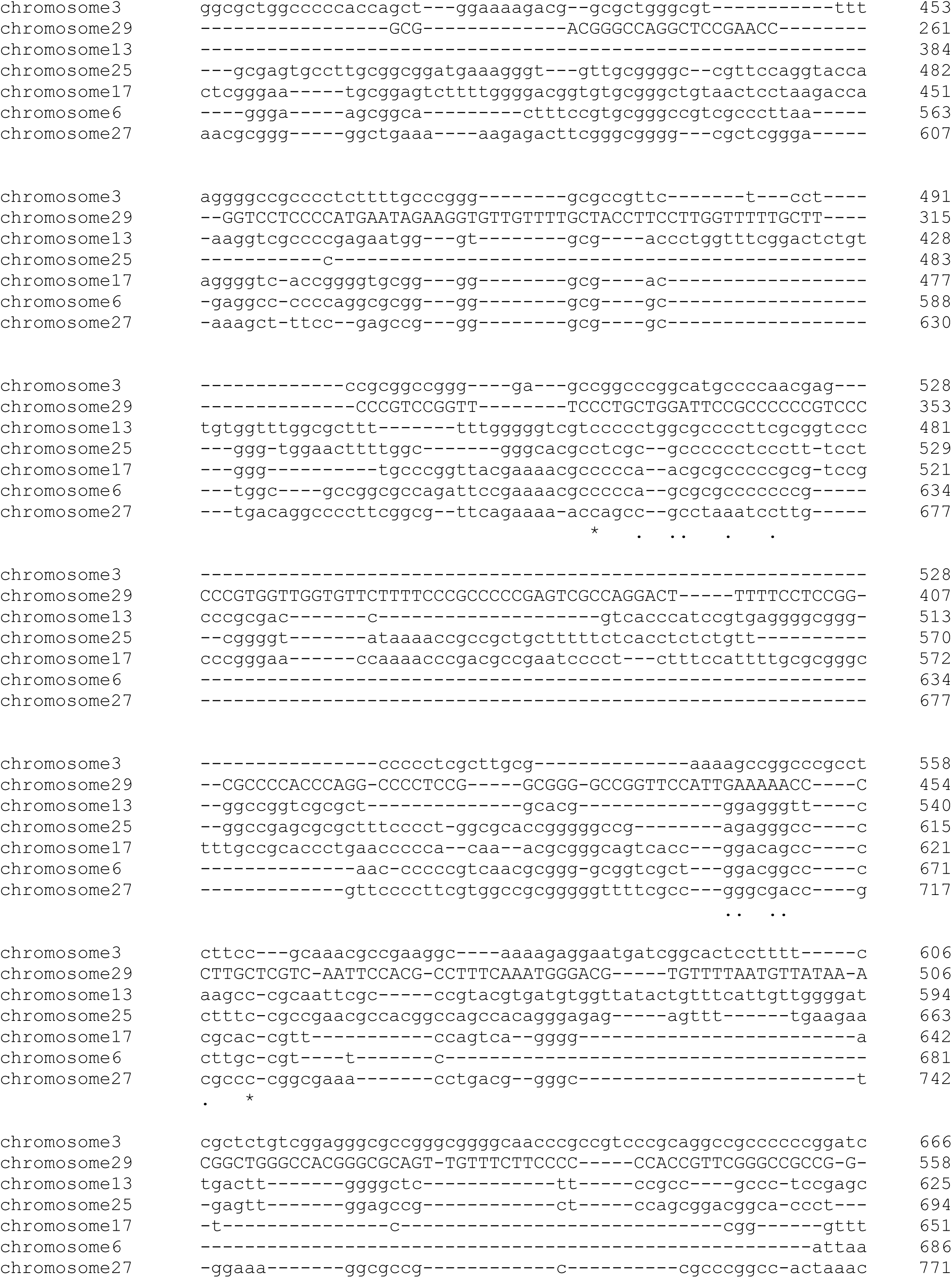
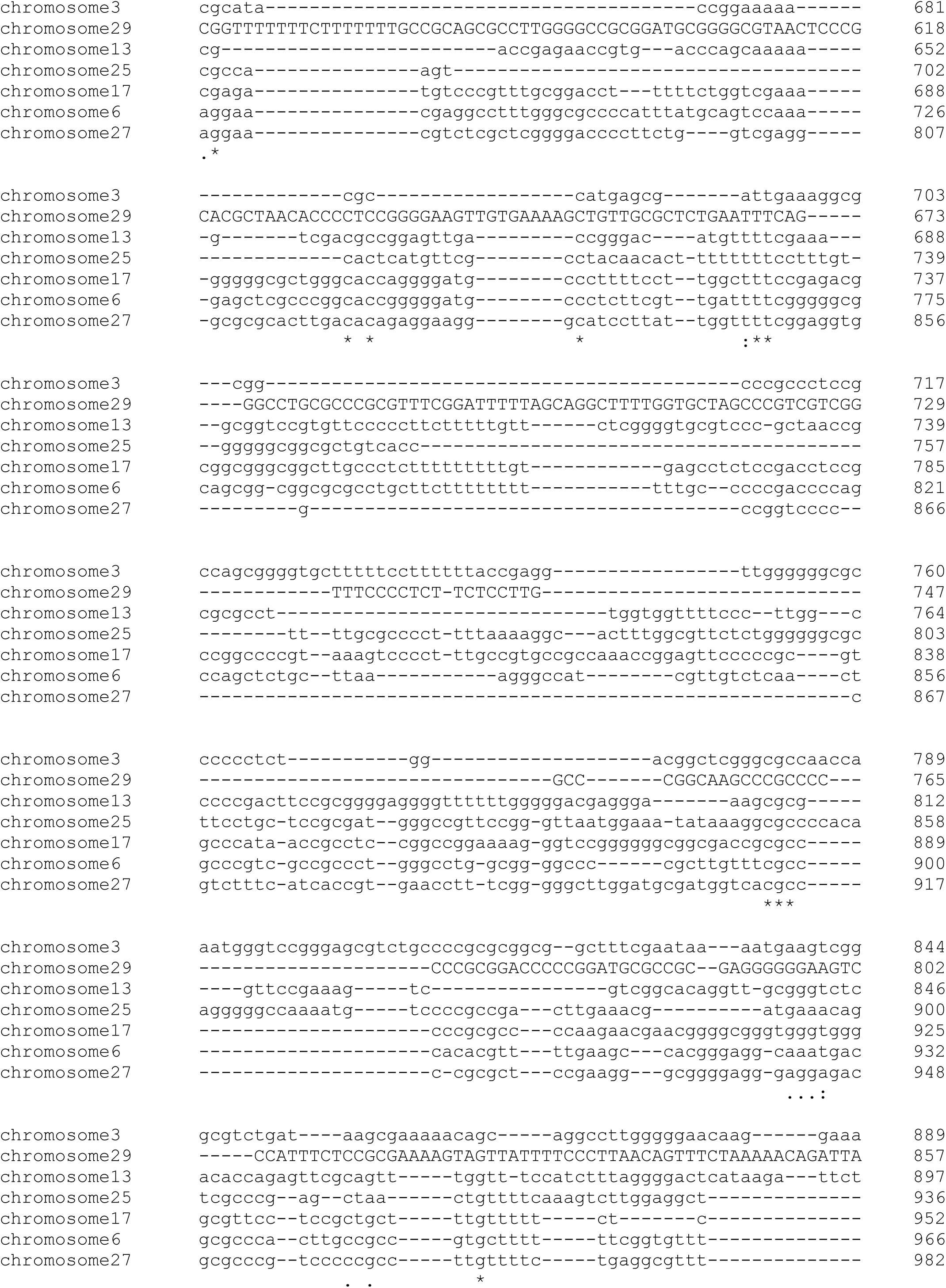
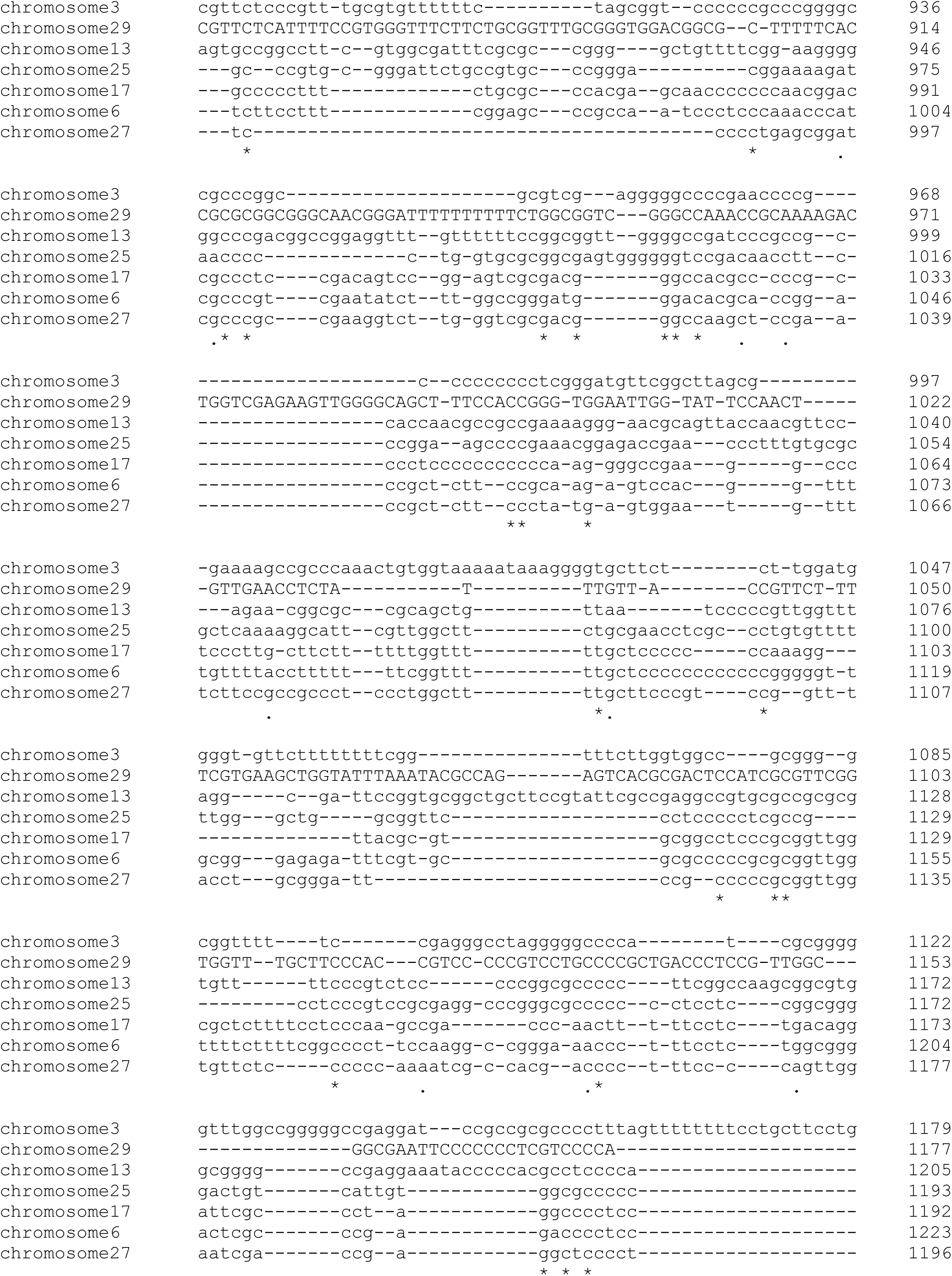
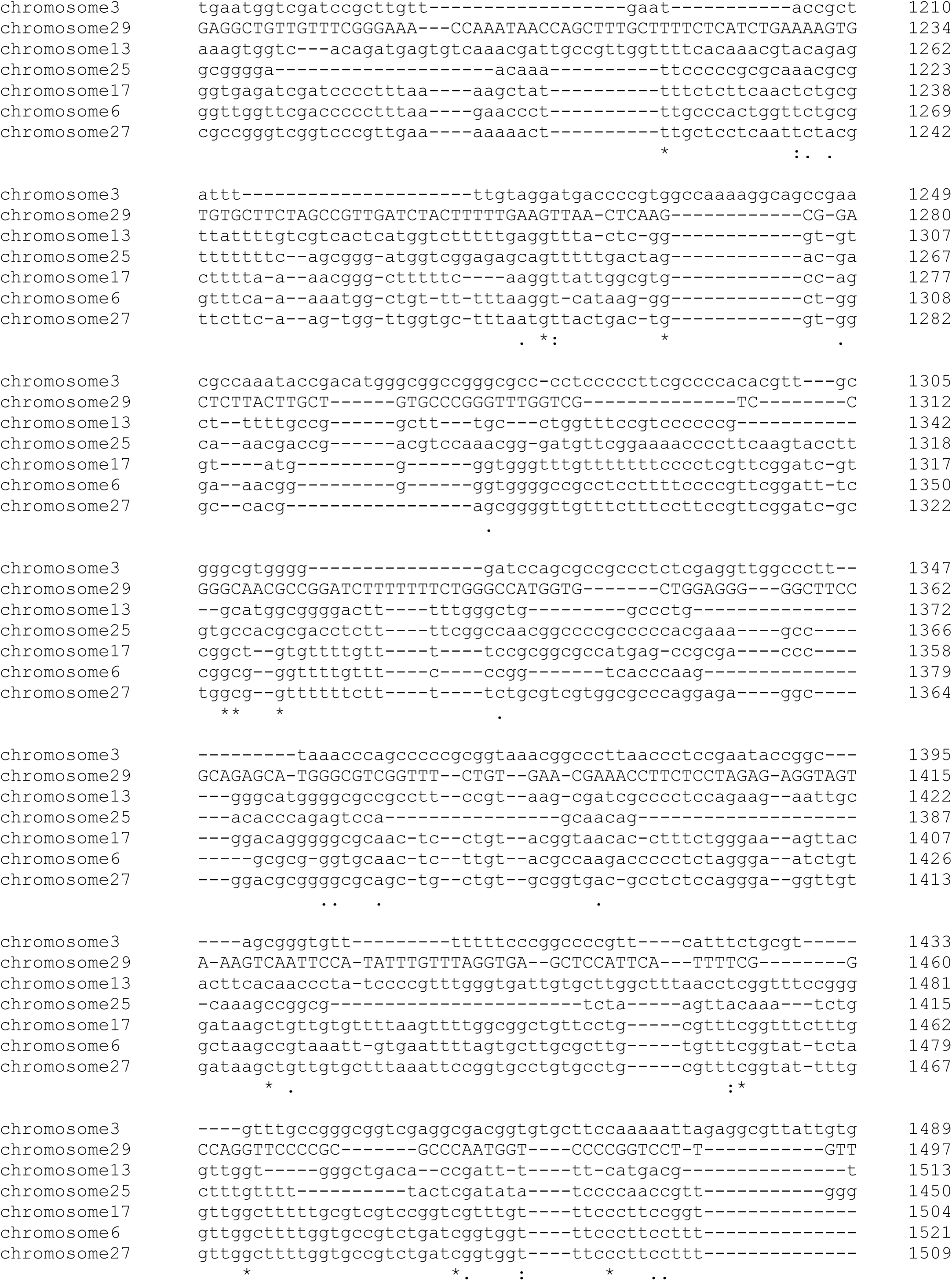
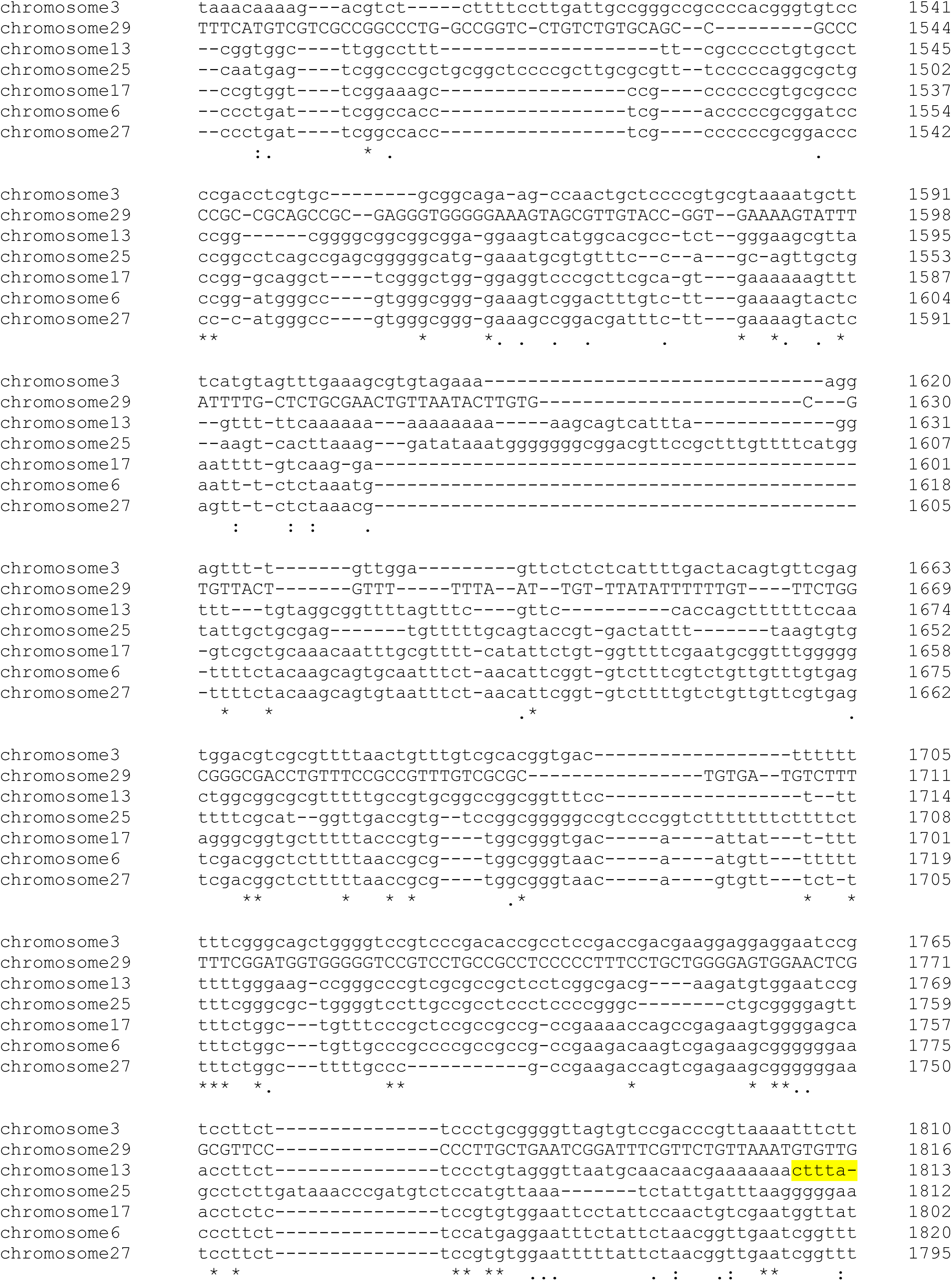
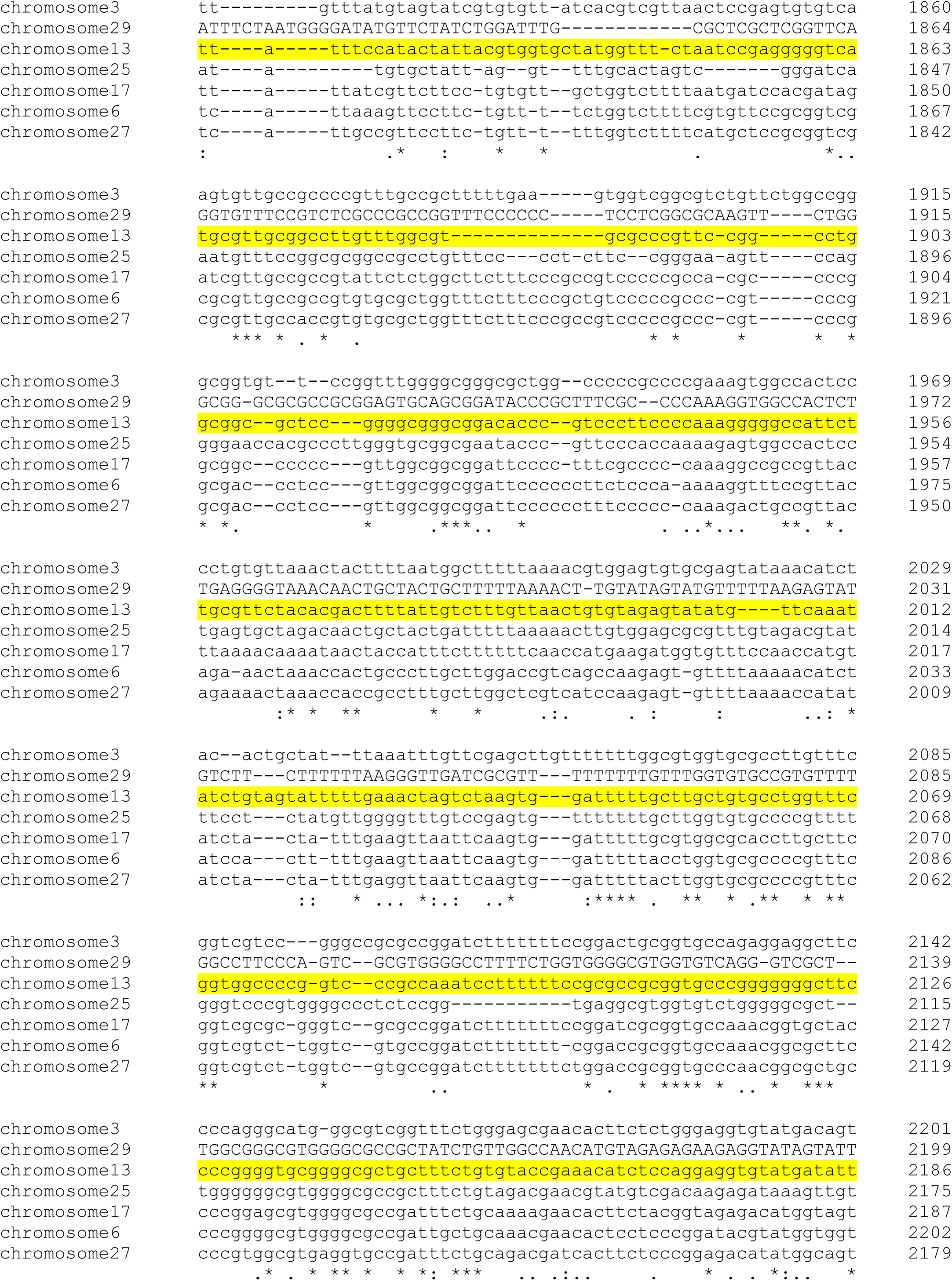
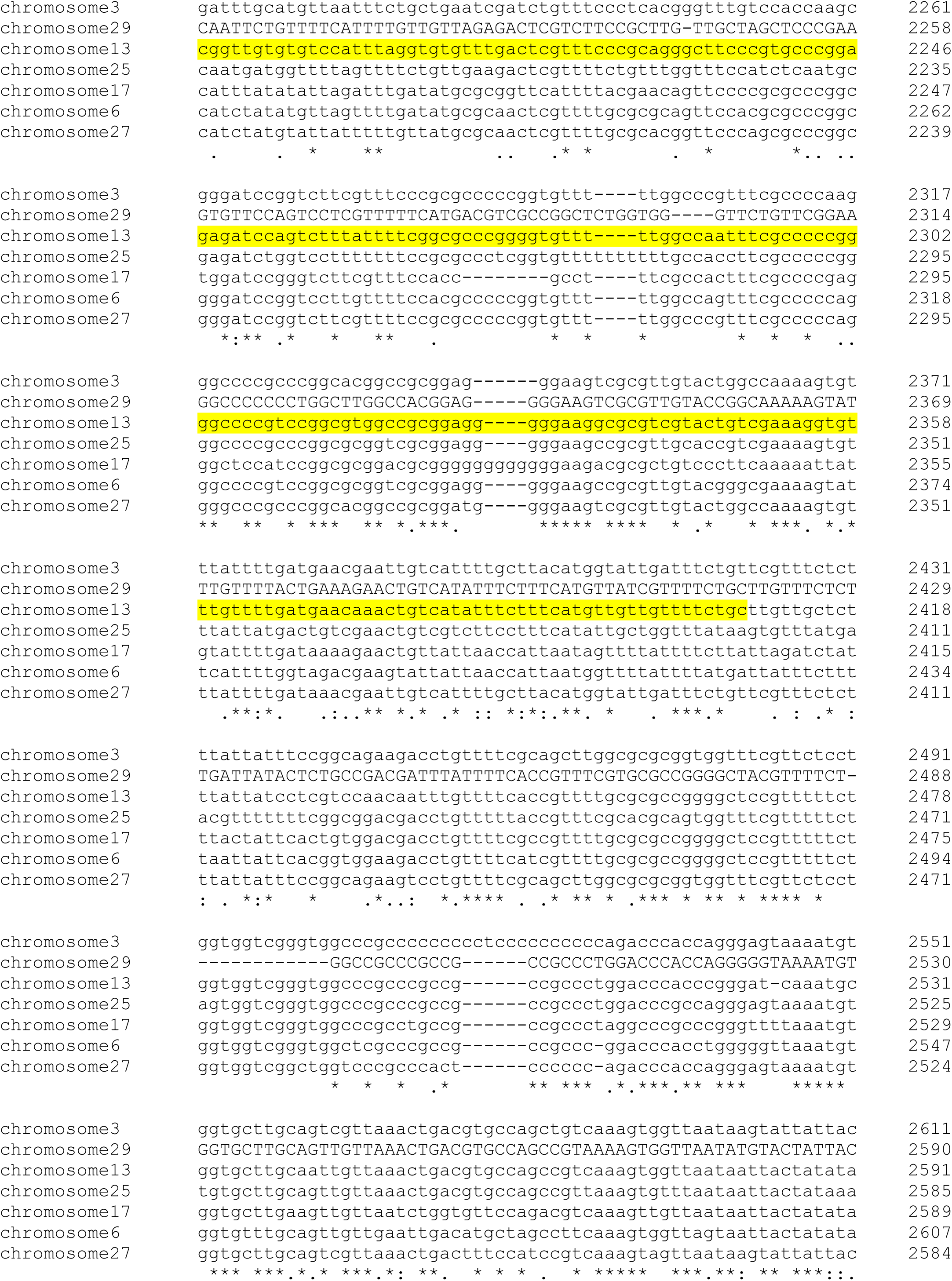
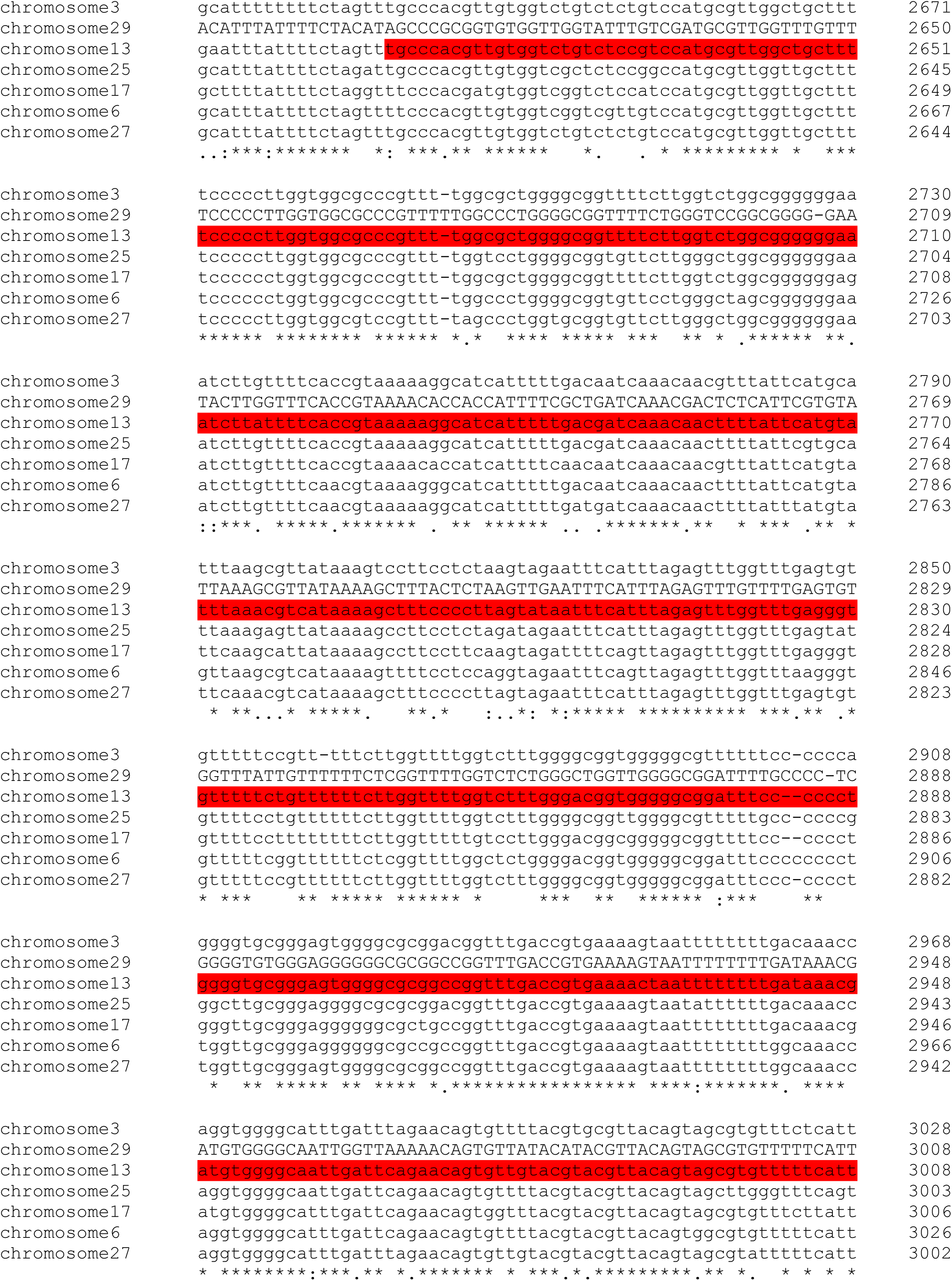
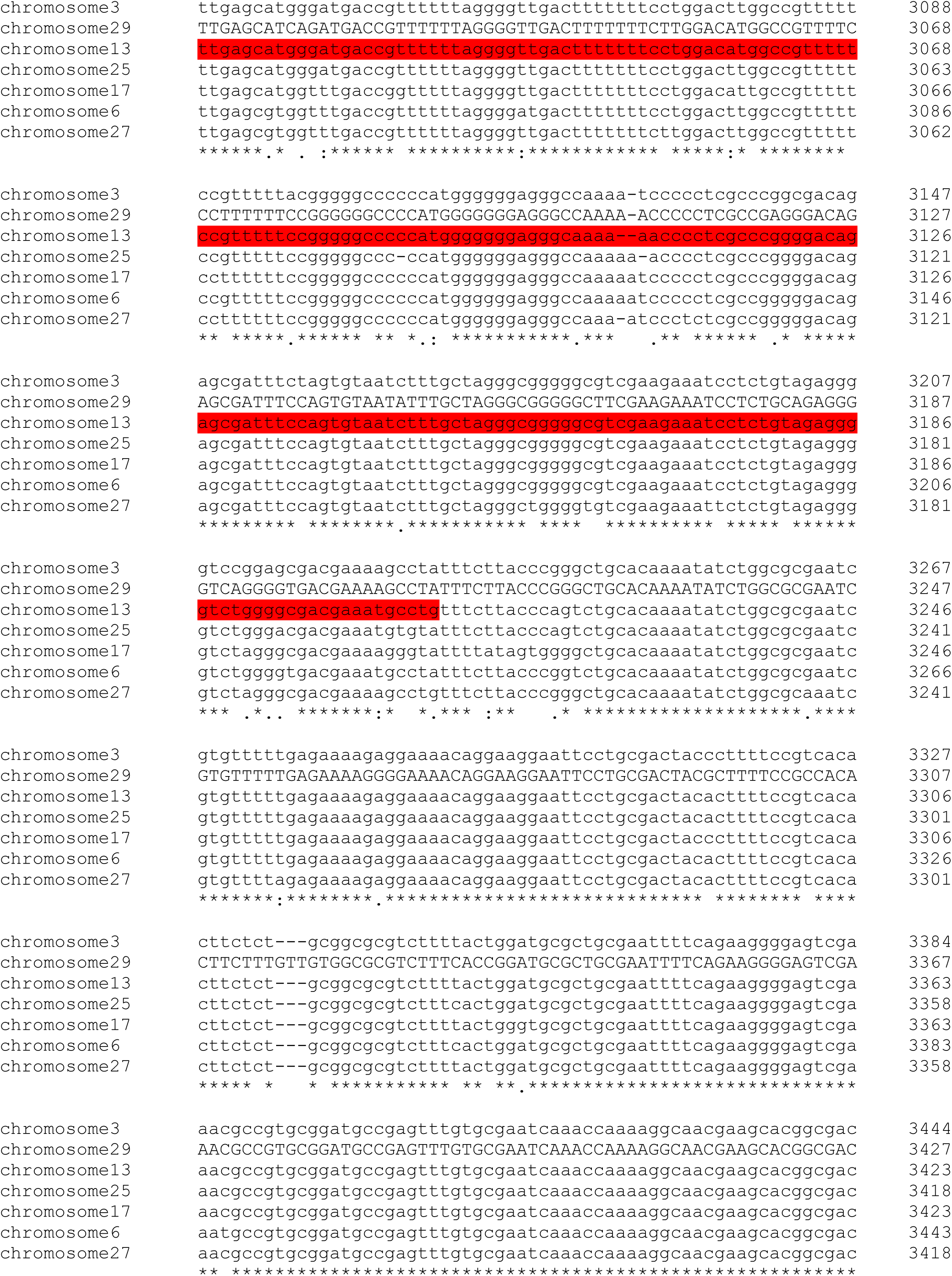
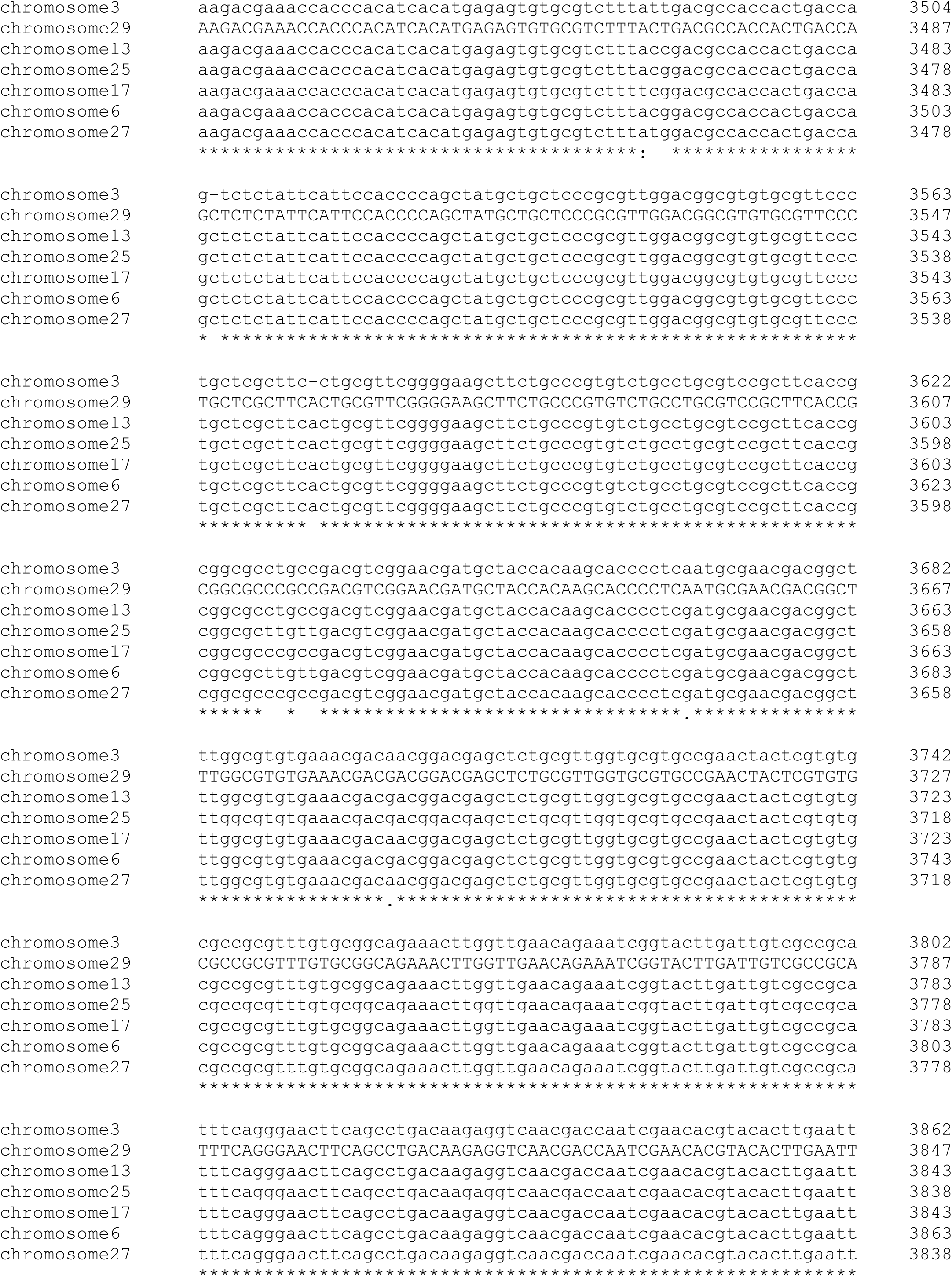
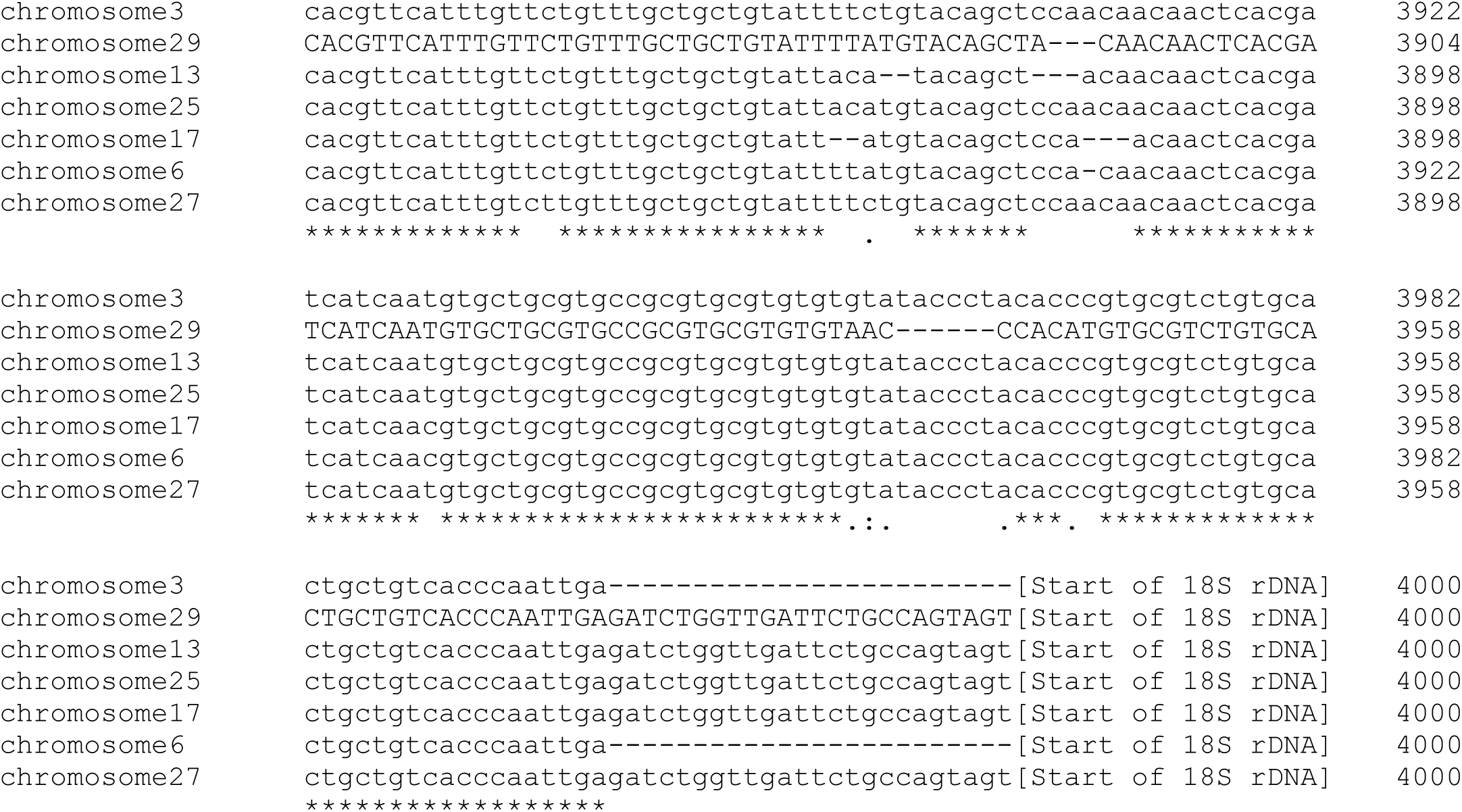
Alignment of the rDNA spacers in *A. deanei*. Regions flanking the insertion site that were used for homologous recombination are highlighted in yellow (3’ flank) and red (5’ flank). rDNA sequences in A. deanei were identified using the rDNA sequence from T. brucei as query (NCBI locus tags TB927_01.rRNA.1 and onwards). For the multi-sequence alignment Clustal Omega was used (www.ebi.ac.uk/jdispatcher/msa/clustalo).

